# Epithelial-to-mesenchymal transition proceeds through directional destabilization of multidimensional attractor

**DOI:** 10.1101/2020.01.27.920371

**Authors:** Weikang Wang, Dante Poe, Yaxuan Yang, Thomas Hyatt, Jianhua Xing

## Abstract

How a cell changes from one stable phenotype to another one is a fundamental problem in developmental and cell biology. Epithelial-to-mesenchymal transition (EMT) is a phenotypic transition process extensively studied recently but mechanistic details remain elusive. Through time-lapse imaging we recorded single cell trajectories of human A549/Vim-RFP cells undergoing EMT induced by different concentrations of TGF-β in a multi-dimensional cell feature space. The trajectories cluster into two distinct groups, indicating that the transition dynamics proceeds through parallel paths. We then reconstructed the reaction coordinates and corresponding pseudo-potentials from the trajectories. The potentials reveal a plausible mechanism for the emergence of the two paths as the original stable epithelial attractor collides with two saddle points sequentially with increased TGF-β concentration, and relaxes to a new one. Functionally the directional saddle-node bifurcation ensures a CPT proceeds towards a specific cell type, as a mechanistic realization of the canalization idea proposed by Waddington.

## INTRODUCTION

Cells of multicellular organisms assume different phenotypes that can have drastically different function, morphology, and gene expression patterns, and can undergo distinct changes when subjected to specific stimuli and microenvironments. Examples of cell phenotypic transitions (CPTs) include cell differentiation during development, induced and spontaneous cell fate transition such as reprogramming and trans-differentiation. Epithelial-to-mesenchymal transition (EMT) is such a prototypic progress that has been extensively studied due to its significance in cell and developmental biology as well as in biomedical research (Fig. 1a). Recent advances in single cell techniques have further accelerated the long-time efforts on unraveling the mechanisms of CPTs, for understanding processes such as differentiation and reprogramming in developmental and cell biology, and for potential biomedical implications of modulating cell phenotypes in regenerative medicine and diseases such as cancer and chronic diseases ^1, 2^.

**Figure 1.**
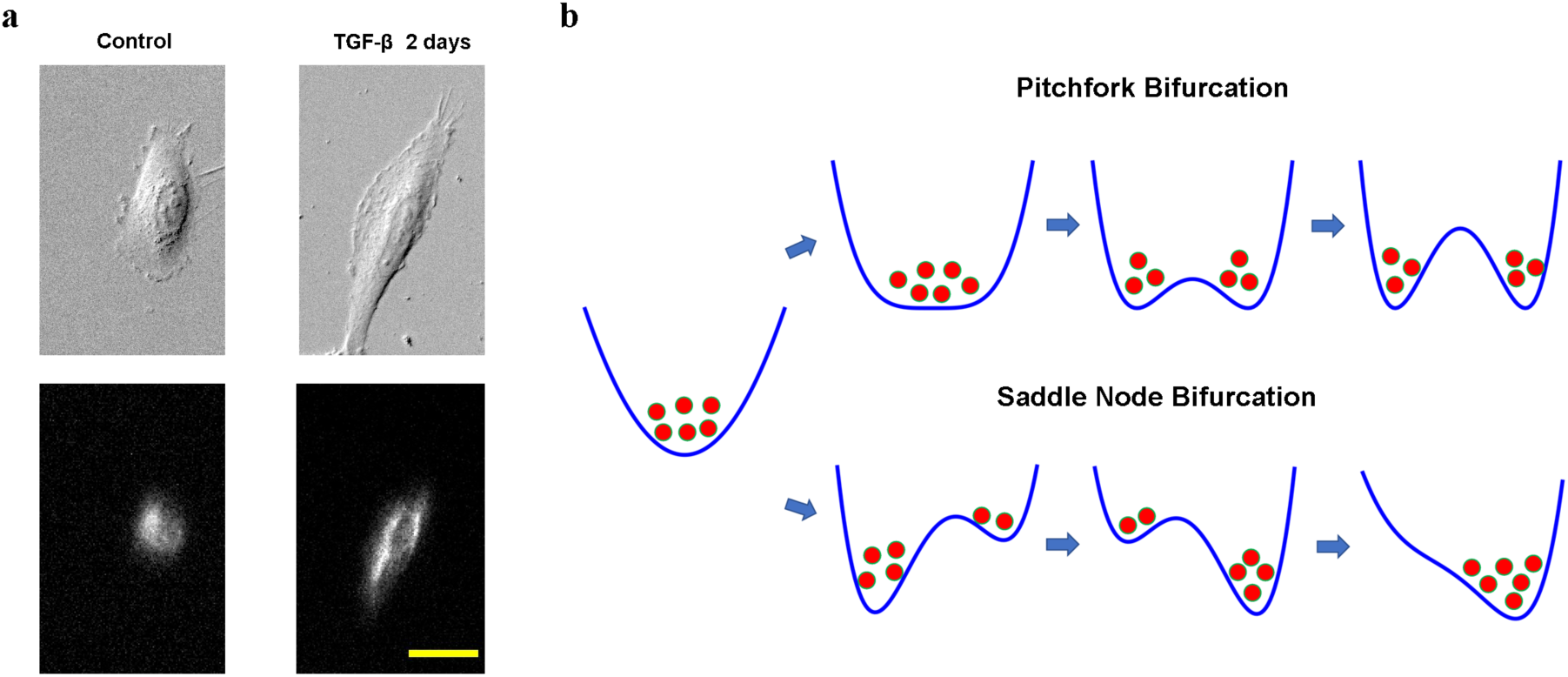
Cell phenotypic transitions as critical state transitions. (a) Transmission light and fluorescence (Vimentin-RFP) images showing an A549/Vim-RFP cell undergoing epithelial-to-mesenchymal transition. Scale bar: 30 µm. (b) Illustration of pitchfork (top) and saddle-node (bottom) bifurcations using 1-D potential systems.

A cell is a dynamical system, and understanding a CPT process from dynamical systems theory is an intriguing long-standing question in mathematical and systems biology. Mathematically at any instant time a cell state can be described by a set of dynamical variables, such as gene expression levels, or other collective cell feature variables. A stable cell phenotype is an attractor in a high-dimensional cell state space formed by the dynamical variables ^3^, and a CPT is a transition between different attractors. Waddington famously made an analogy of developmental processes to a ball sliding down a potential landscape, and diverging into multiple paths at some bifurcation points. This picture exemplifies a pitchfork bifurcation, where an original stable attractor turns to an unstable saddle point together with emergence of two new attractors (Fig. 1b top). Another well-discussed mechanism for attractor-to-attractor transition is a saddle-node bifurcation, where first a new attractor emerges, and the saddle point separating the two attractors moves towards to and eventually collides with the old attractor to destabilize the latter (Fig 1b bottom). A notable difference between the two types of bifurcation is that after pitchfork bifurcation the original state can relax to multiple new stable states, while for the saddle-node bifurcation the system is directed to relax to one attractor. It is a fundamental theoretical question which mechanism a CPT process assumes, and why.

The pitchfork bifurcation and saddle-node bifurcation are two theoretical mechanisms of critical state transition that have been discussed in the context of CPTs ^4^. A new challenge then is how to evaluate various mathematical models experimentally at single cell levels. While several studies have suggested such critical state transitions through analyzing snapshot single cell genomics data ^4, 5^, the fixed cell data cannot provide temporal information on how individual cells transit. On the other hand, information from fluorescence-based live cell imaging is typically restricted to a small number of molecular species. Tracking a small number of molecular species through live cell imaging cannot provide the collective transition dynamics due to the intrinsic high-dimensional nature.

Specific for EMT, theoretical studies suggest it proceeds as saddle-node bifurcation^6^, but there has no direct experimental study on the single cell transition dynamics. As recognized in a consensus statement from researchers in the EMT field, several open questions and challenges exist on mechanistic understanding of the EMT process ^7^. For example, it is unclear whether the process proceeds as hopping among a small number of discrete and distinct intermediate states, or a continuum of such states with no clear boundary. The transition may proceed either along a linear array of the states or through multiple parallel paths. Pseudo-time analyses of high throughput single cell genomics studies infer that EMT proceeds through a 1-D continuum path ^8^, consistent with a prevalent EMT axis concept with the epithelial and the mesenchymal states as the two end states ^9^. These predictions, which are indirectly inferred from these snapshot single data, require direct test through tracking single cells over time, but live cell imaging studies are impeded by the observation that the EMT status cannot be assessed based only a small number of molecular markers including regulatory elements such as key transcriptional factors ^7^.

To address the above challenge on studying EMT dynamics, recently we developed a platform of tracking cell state change in a composite cell feature space that is accessible for multiplex and long-term live cell imaging ^10^. Here we first apply the platform to study EMT in a human A549 derivative cell line with endogenous vimentin-RFP labeling (ATCC^®^ CCL-185EMT^™^, denoted as A549/Vim-RFP in later discussions) induced by different concentrations of TGF-β. Then we analyze an ensemble of recorded multi-dimensional single cell trajectories within the framework of reaction rate theories that have been a focused subject in the context of physics and chemistry.

## RESULTS

### Single cell trajectories are mathematically represented in a collective morphology/texture feature space

For quantitative studies of CPT dynamics, first one needs to establish a mathematical representation of the cell states and cell trajectories. Mathematically one can represent a cell state by a point in a multi-dimensional space defined by gene expression ^11^ or other cell properties ^12^. Noticing that cells have phenotype-specific morphological features that can be monitored even with transmission light microscopes, recently we developed a framework that defines cell states in a combined morphology and texture feature space ^10^. The framework allows one to trace individual cell trajectories during a CPT process through live-cell imaging. We applied the framework to study the TGF-β induced EMT with the A549/Vim-RFP cells (Fig. 2) ^10^. Vimentin is a type of intermediate filament protein commonly used as a mesenchymal marker, and changes its expression and spatial distribution during EMT ^13^. Furthermore, during EMT cells undergo dramatic change of cell morphology accompanying gene expression change (Fig. 1a) ^13, 14^. Accordingly, we used a combination of vimentin texture and cell shape features to specify a cell state.

**Figure 2.**
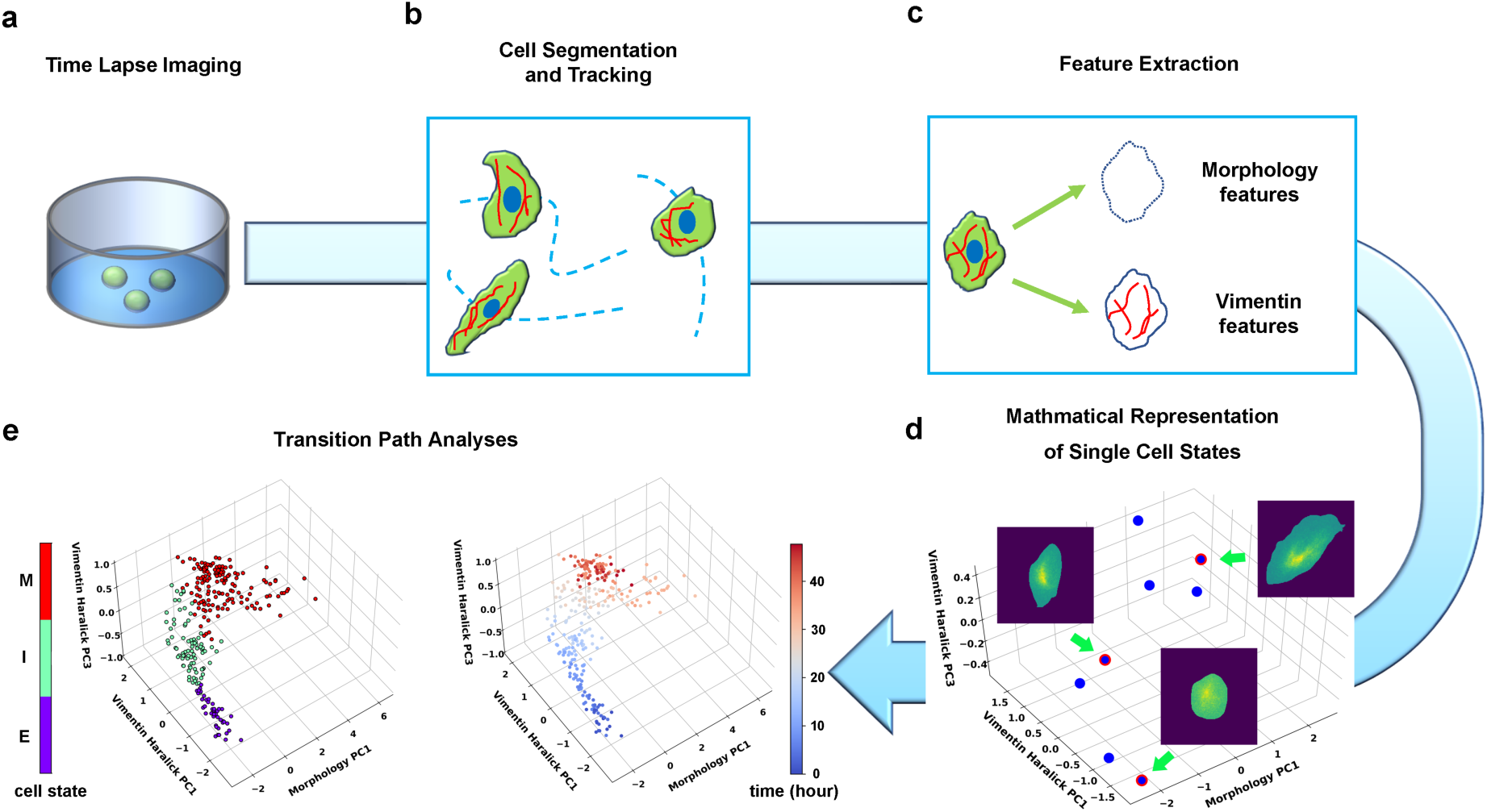
Summary of pipeline for recording and analyzing single cell trajectories in composite multi-dimensional cell feature space. (a) Time lapse imaging of A549/Vim-RFP cells treated with TGF-β. (**b**) Deep-learning aided single cell segmentation and tracking on the acquired time lapse images. (**c**) Extraction of morphology and vimentin features of single cells. (**d**) Representation of single cell states in a multidimensional morphology/texture feature space. (**e**) Transition path analyses over recorded trajectories. Right: A representative single cell trajectory of EMT in the feature space. Color represents time (unit: hour). Left: the same trajectory colored by the regions in the feature space (E, I, and M, for epithelial, intermediate, and mesenchymal regions, respectively) each data point resides. Reduced units are used in this and all other figures.

We first performed time lapse imaging of the EMT process of A549/Vim-RFP induced with 4 ng/ml TGF-β (Fig. 2a, Methods (1)), then performed single cell segmentation and tracking on the acquired images (Fig. 2b). We quantified the images with an active shape model ^15^ and performed principal component analysis (PCA) to form a set of orthonormal basis vectors of collective variables, which include cell body shape of 296 degrees of freedom (DoF) quantified by an active shape model ^16^, and texture features of cellular vimentin distribution quantified by 13 Haralick features ^17^ (Fig. 2c). Then the state of a cell at a given instant is represented as a point in the composite morphology/texture feature space (Fig. 2d), and the temporal evolution of the state forms a continuous trajectory in the space subject to further theoretical analyses (Fig. 2e).

### Transition path analyses identify an ensemble of reactive single cell trajectories

Before TGF-β treatment, a population of cells assumes a localized stationary distribution in this 309-dimensional composite space, and most cells are epithelial (Fig. 3a, blue). TGF-β treatment destabilizes such distribution, and the cells relax into a new stationary distribution dominated by mesenchymal cells (Fig. 3a red). We recorded 204 continuous trajectories in the state space. A representative trajectory shown in Fig. 2e (right) (also in Movie. S1) reveals how a cell transits step-by-step from an epithelial cell with round polygon shapes and a localized vimentin distribution, to the mesenchymal phenotype with elongated spear shapes and a dispersive vimentin distribution.

**Figure 3.**
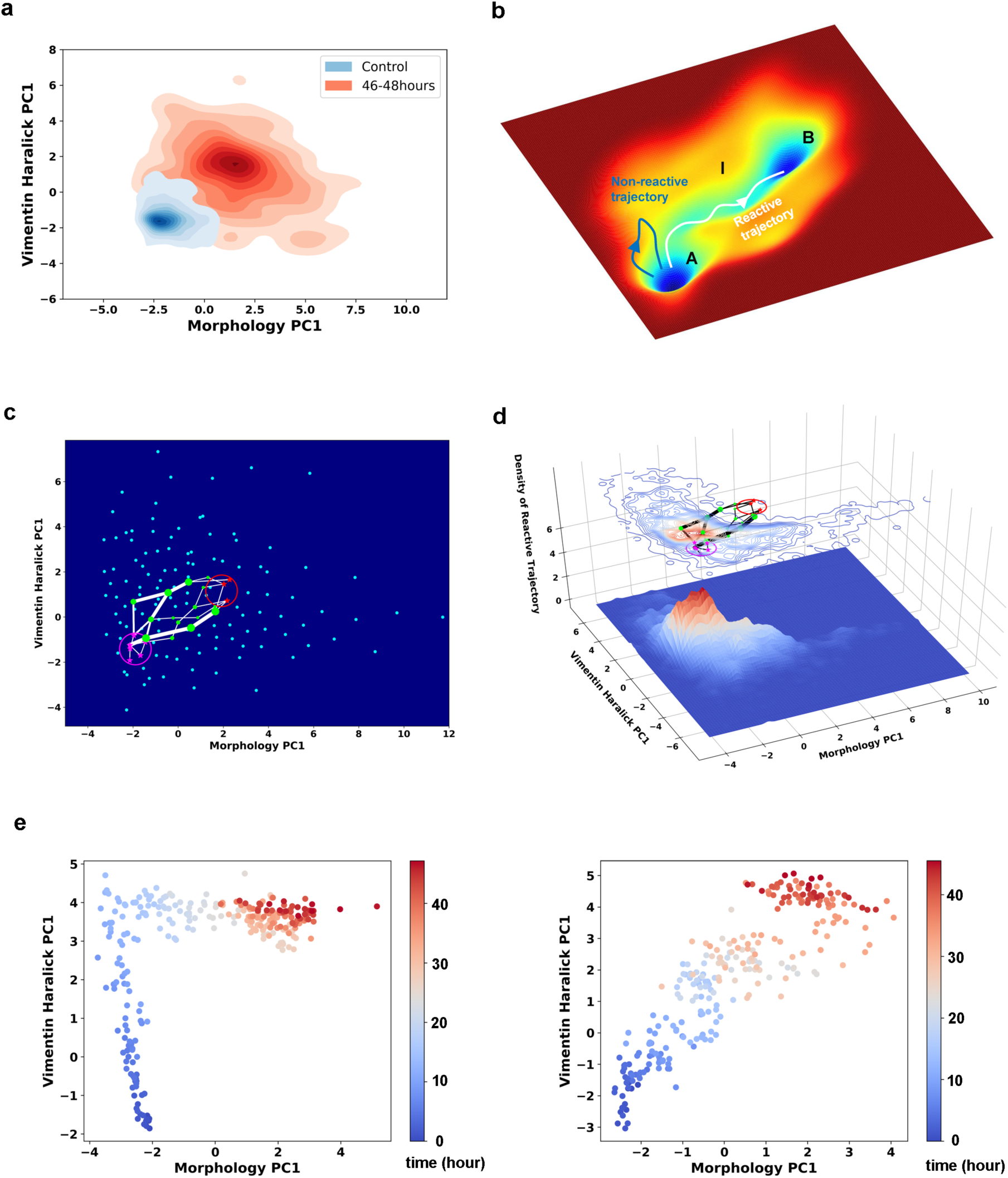
Single cell trajectories of EMT form two distinct groups. **(a)** Kernel density estimation of cells in control condition (Blue) and cells after treated with 4ng/ml TGF-β for 46-48 hours (Red). **(b)** Example reactive and nonreactive trajectories in a quasi-potential system. Also shown is a valley (tube) connecting regions A and B that most reactive trajectories fall in. **(c)** Space approximation of the whole single cell data set with self-organizing map into 12×12 discrete states (clusters). Directed network generated base on the self-organizing map and the transition between states. The distance between two states is defined as the negative logarithm of transition probability. Shortest paths (white lines) between epithelial states (purple stars) and mesenchymal states (red stars). Green dots are the states that the shorted paths passed by. The size of a dot stands for the frequency of this dot passed by shortest paths. The width of a white line represents the frequency that these shortest paths passed by. **(d)** Contour map (top, superimposed with the shortest paths in panel **c**) and 3D surface-plot (bottom) of density of reactive trajectories in the plane of morphology PC1 and vimentin Haralick PC1. **(e)** Representative trajectories from the two groups. Left: Vimentin varies first. Right: Concerted variation.

Next, we applied rate theory analyses on the recorded single cell trajectories. Rate theories study how a system escapes from a metastable state, or relaxation from one stationary distribution to a new one ^18^. Specifically a modern development in the field is the transition path theory (TPT). Within the TPT framework, one divides the state space into regions containing the initial (*A*) and final (*B*) attractors, and an intermediate (*I*) region. A reactive trajectory is one that originates from region *A*, and enters region *I* then *B* before re-entering region *A* (Fig. 3b). The reactive trajectories form an ensemble of transition paths that connect regions *A* and *B*.

Accordingly, we divided the four-dimensional principal component subspace into epithelial (*E*), intermediate (*I*), and mesenchymal (*M*) regions (Fig. 2e left) ^10^. With this division of space we identified a subgroup of 135 recorded single cell trajectories that form an ensemble of reactive trajectories that connect *E* and *M* by day 2. It should be noted that the definition of reactive trajectories here differs from that in the classical transition path theories, which refer segments of trajectories of infinite length that travel back and forth between the initial and final regions numerous times.

### TGF-β induced EMT in A549/Vim-RFP cells proceed through parallel transition paths

We first examined whether EMT proceeds through one or multiple types of paths through analyzing the ensemble of reactive trajectories using self-organizing map (SOM) (Methods (2)). SOM is an unsupervised artificial neural network that utilize neighborhood function to represent the topology structure of input data ^19^. The algorithm clusters the recorded cell states into 144 discrete states, and represents the EMT process as a Markovian transition process among these states (Fig. S1). Shortest path analysis using the Dijkstra algorithm ^20^ over the transition matrix reveals two groups of paths: vimentin Haralick PC1 varies first, and concerted variation of morphology PC1 and vimentin Haralick PC1, with finite probabilities of transition between the two groups (Fig. 3c). This result is consistent with our previous trajectory clustering analysis using dynamics time warping (DTW) distance between reactive trajectories ^10^. To further support this conclusion, we examined the density of reactive trajectories ρ_R_, in the plane of morphology PC1 and vimentin Haralick PC1 (Methods (3)). The contour map of ρ_R_ shows two peaks corresponding of the two groups of shorted paths in the directed network (Fig. 3d). The peak that vimentin Haralick PC1 varies first is higher than the peak of concerted variation, indicating more reactive trajectories along this path. The distinct features of the two types of trajectories are apparent from the two representative single cell trajectories shown in Fig. 3e.

### A revised string method is developed to reconstruct reaction coordinate from single cell trajectories

Theoretically, for a stochastic system, one can define an action *S, e.g*., the Onsager-Machlup action for a reactive trajectory, so the probability of observing this trajectory is exp (−*S*). Reactive trajectories mainly concentrate within a tube in the state space that connects regions *A* and *B* (Fig. 3b). The center of the tube has been used to define a reaction coordinate (RC), a concept central to rate theories (Fig. 4a) ^21^. A RC is a one-dimensional geometric parameter (denoted as *s* in the subsequent discussions) that describes the progression along a continuous reaction path defined in the state space ^22^. A good choice of RC can provide mechanistic insight on how the transition process proceeds.

**Figure 4.**
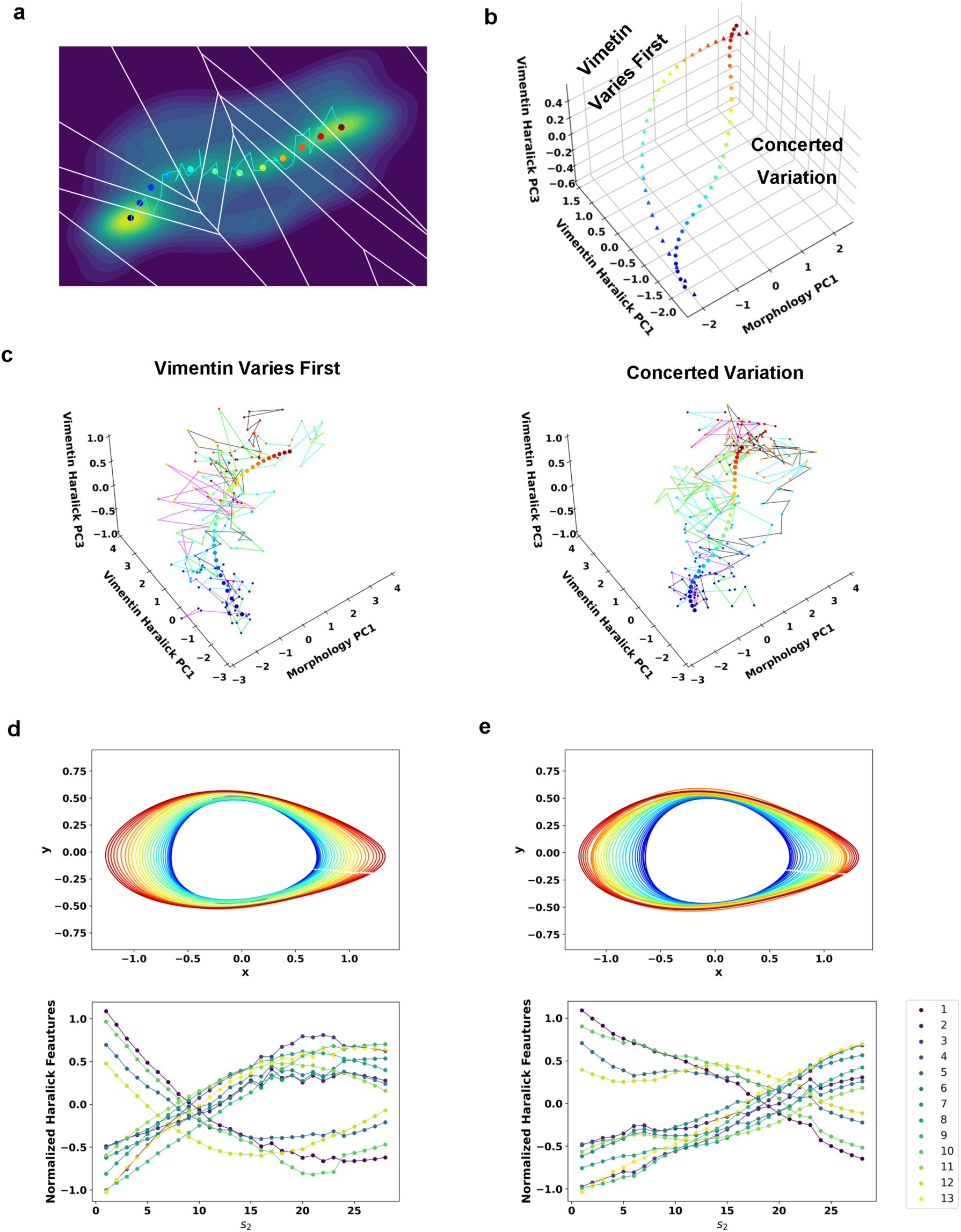
Reaction coordinate reconstruction of two parallel paths from reactive A549/Vim-RFP EMT trajectory ensemble with a modified string method. **(a)** Discrete representation of a 1-D reaction coordinate (colored dots) on the filled contour map with corresponding Voronoi cells. The cyan line is a reactive trajectory that starts from A and ends in B. **(b)** Reconstructed RCs from reactive single cell EMT trajectories using a revised finite temperature string method. **(c)** Separate representation of the RCs overlaid with representative reactive single cell trajectories. Left: vimentin varies first; Right: concerted variation. **(d)** Cell shape (top), Haralick feature (bottom) along RC s_1_. The colors of the cell shapes in b/c left represent progression of EMT (starts from blue and ends in red). Haralick feature 1: Angular Second Moment; 2: Contrast; 3: Correlation; 4: Sum of Squares: Variance; 5: Inverse Difference Moment; 6: Sum Average; 7: Sum Variance; 8: Sum Entropy; 9: Entropy; 10: Difference Variance; 11: Difference Entropy; 12: Information Measure of Correlation 1; 13: Information Measure of Correlation 2. **(e)** Similar to panel e but along RC s_2_.

Therefore, we set to reconstruct the RC for each path with the recorded reactive trajectories using a revised finite temperature string method ^23, 24, 25^. The method first approximates the RC by a set of discrete image points ^26^ that are uniformly distributed along the arc length of the RC, so the multi-dimensional state space can be divided by a 1-D array of Voronoi polyhedra containing individual images, and the data points can be assigned to individual Voronoi cells (Fig. 4a). Starting with a trial RC to define the initial division of the Voronoi cells, one optimizes the trial RC iteratively by minimizing the distance dispersion between the string point and sample points within each Voronoi cell. Since here we have an ensemble of continuous trajectories, we modified the iteration procedure slightly. Specifically, we minimized both the distance between the ensemble of measured reactive trajectories and the image point within each individual Voronoi cell, as well as the overall distance between each individual trajectory and the trial RC (Fig. S2, Methods (4)).

Therefore, we grouped the reactive trajectories based on the DTW distance, then identified the RCs for each group separately following the modified string method. The iteration procedure gives the RC of each path (*s*_1_ and *s*_2_) that characterizes common features of the reactive trajectories for the TGF-β induced EMT in A549/Vim-RFP cells (Fig. 4b). The recorded trajectories fluctuate around the RCs and form reaction tubes as expected from the TPT theory (Fig. 4c). Along each RC the cell shape changes dominantly through elongation and growth (Fig. 4d & e top), and most of the 13 vimentin Haralick features increase or decrease monotonically and continuously over time (Fig. 4d & e, bottom) (Methods (5)). The two RCs first diverge from the *E* region to follow two distinct paths, then converge within the *M* region. In one group (Fig. 4d), most of the Haralick feature changes take place before major morphology change. For the group with concerted dynamics (Fig. 4e), both cell shape and Haralick features vary gradually along the RC.

### Reconstructed reaction coordinate and pseudo-potentials reveal EMT as relaxation along continuum manifolds

Considering a cell as a dynamical system, with the single cell trajectories one may formulate an inverse problem to reconstruct the underlying dynamical equations governing the EMT transition. Take a minimal ansatz that assumes the dynamics of the collective variables (**x**) can be described by a set of Langevin equations in the morphology/texture feature space, *d***x** / *dt* = **F**(**x**) + *η*(*t*), where **F**(**x**) is a governing vector field, and *η* are white Gaussian noises with zero mean. Then in principle one can reconstruct **F**(**x**) from the single cell trajectory data, **F**(**x**) = ⟨*d***x** */ dt*⟩, averaged over the neighborhood of each **x**. Notice for the ensemble average one needs to use all the reactive and nonreactive trajectories.

For mathematical simplicity and better numerical convergence, here we restricted to reconstructing the dynamics along the RC (Fig. 5a). The ansatz becomes a 1-D convection-diffusion process, *ds* / *dt* = −*dϕ* / *ds* +*η*. Notice that for a 1-D system even without detailed balance one can define a pseudo-scalar potential *ϕ* ^27, 28^. This pseudopotential corresponds to a potential of mean force in the case of a conserved system. To confirm that the ansatz of using a memory-less 1-D Langevin equation is a good approximation for the EMT dynamics along a RC, we performed the Chapman-Kolmogorov Test (CK-test). The test shows that the 1-step transition matrix can indeed predict the dynamics on longer time scales, and thus the Markovian assumption is a good zero^th^-order approximation of the EMT dynamics (Fig. S3, Methods (6)).

**Figure 5.**
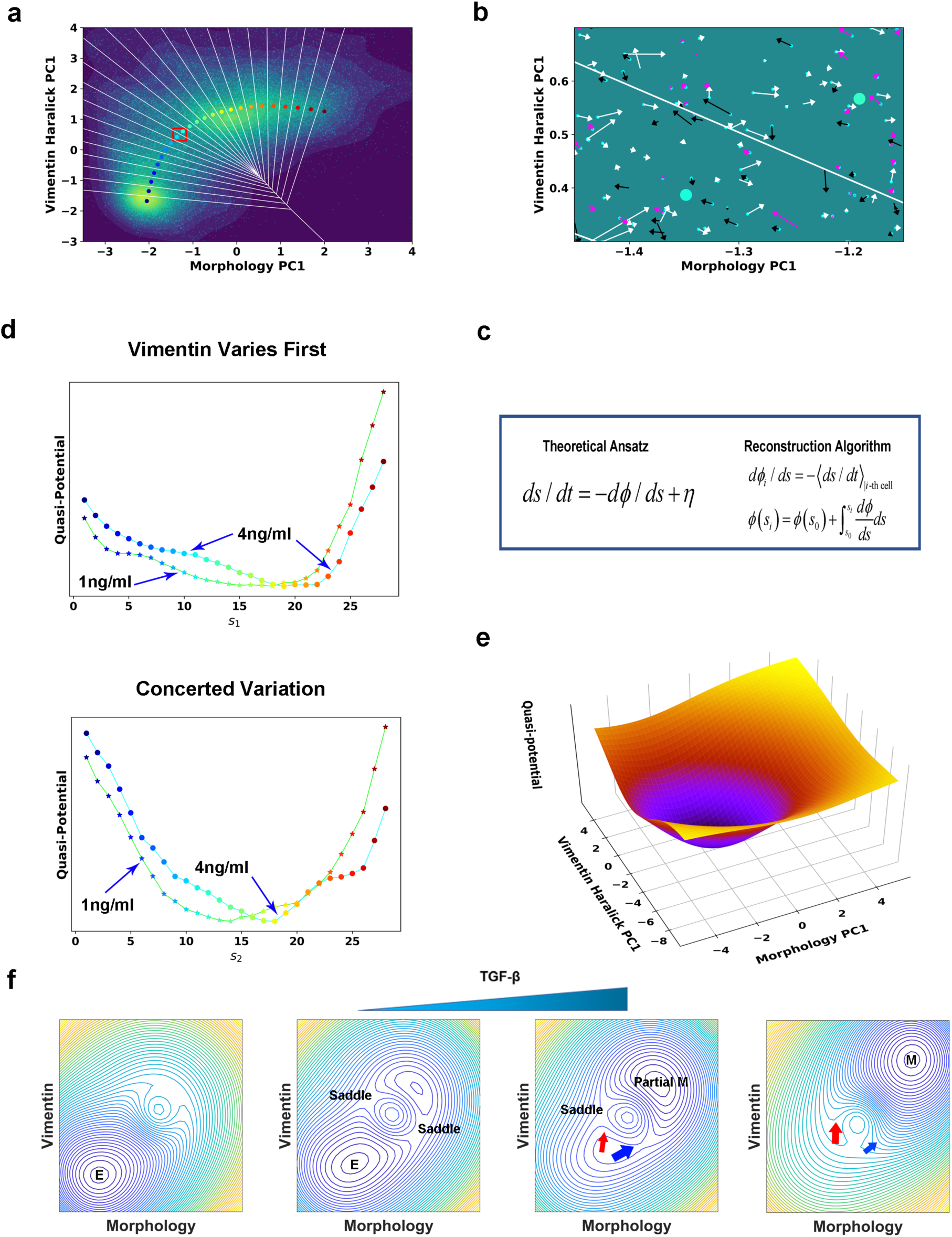
Quantification of dynamics along the two RCs suggest a mechanism of forming two EMT paths through sequential saddle-node collisions. (**a**) Reconstructed RC s_1_ on density plot of single cell trajectories data. Cyan dots are single cell data points. (**b**) Enlarged view of the red box region in Fig. 5a. The arrow associated with each data point (cyan dot) represents the value of ds/dt (white: >0, magenta: = 0, black: <0). (**c**) Theoretical framework of dynamics reconstruction along the RC. (**d**) Comparison of reconstructed pseudo-potentials along the RC with 1ng/ml and 4ng/ml TGF-β treatment. star (lime color line): 1ng/ml; circle (cyan color line): 4ng/ml. Top: Reconstructed pseudo-potentials along RC s_1_. Bottom: Reconstructed pseudo-potentials along RC s_2_. (**e**) Pseudo-potential of control situation based on kernel density estimation. (**f**) A metaphorical potential system to illustrate a plausible mechanism for generating the two paths through sequential collision between a stable attractor and two saddle points when the concentration of TGF-β increases. The width of the arrows represents the probabilities that single cell trajectories follow corresponding paths. E:epithelial attractor; Partial M: partial EMT attractor; M: mesenchymal attractor.

To reconstruct the dynamical equations, we used the measured trajectories and instant velocities (along the RC direction) as input. An enlarged view in Fig. 5b shows the instant velocities (*ds/dt*) of various trajectories segments within Voronoi cells. Numerically we related the potential gradient *dϕ/ds* with the mean velocity within the *i*_*th*_ Voronoi cell through averaging over all recorded trajectory segments that locate within the Voronoi cell at any time *t* (Fig. 5a & c, Methods (7, 8)). On the obtained curve of *dϕ/ds* vs. *s* (Fig. S4), the zeroes correspond to stationary points of the potential. We then reconstructed the pseudo-potential through integrating over *s*, 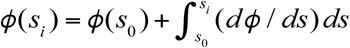 (Fig. 5d). We also obtained the pseudo-potential of the untreated cells (Fig. 5e) from the steady state distribution of untreated cells along the RC, *ϕ*_0_ ∝ − log *p*_*ss*_.

The reconstructed RCs and pseudo-potentials provide mechanistic insight on the transition. Before TGF-β induction, cells reside on the untreated cell potential centered with a potential well corresponding to the epithelial attractor (Fig. 5e). After induction, the system relaxes following the new potential into a new well corresponding to the mesenchymal attractor. Notably in the new potential the original epithelial attractor disappears, reflecting that the epithelial phenotype is destabilized under the applied 4 ng/ml TGF-β concentration.

With the reconstructed pseudo-potentials and variance of *dϕ/ds* obtained from experiment, we solved the Fokker-Planck (FP) equation corresponding to the Langevin equation to predict the steady distribution vs. RC (Methods (9)). While the dynamical equations were reconstructed from trajectories during the transition process and with a minimal Langevin equation ansatz, the predicted stationary distribution matches reasonably well with the distributions sampled from the final stage trajectory data, i.e., after 39 h TGF treatment (Fig. S5 a&b).

### Lowering TGF-β concentration leads to relaxation to a new partial EMT attractor through two parallel paths

Notice that pseudopotential with 4 ng/ml TGF-β for the Vimentin varies first path shows a plateau around s_1_ = 8 (Fig. 5d). We hypothesized that this plateau is a remnant of the original epithelial attractor, and if so we expect to observe a flatter plateau or even a metastable attractor with a lower TGF-β concentration. Therefore we treated A549/Vim-RFP cells with 1 ng/ml TGF-β and recorded 135 three-day long single cell trajectories. Among the recorded trajectories 65 indeed reached the *M* region. The reactive trajectories also cluster into two groups (Fig. S6). For the purpose of comparison we projected the trajectories onto the RCs obtained from the 4 ng/ml TGF-β and reconstructed the pseudo-potentials in the case of 1 ng/ml of TGF-β. Indeed the pseudo-potential for the path with vimentin varying first has a plateau flatter than that of the 4 ng/ml TGF-β treated cells. The pseudopotential of another path, however, does not show such flattening.

In the EMT field it is under debate whether EMT has a discrete or continuum spectrum of intermediate states. If EMT proceeds through a set of discrete and definite states, we expect to observe attractors, i.e., potential wells, corresponding to these states, and lowering TGF-β concentration leads to increased barrier between neighboring wells and cells trapped at some intermediate states for long time. While all the pseudo-potentials at the two TGF-β concentrations reveal destabilization of the epithelial attractor, they do not have multiple attractors anticipated for discrete EMT states (Fig. 5e, *dϕ/ds* in Fig. S7 a&b). Compared to those at 4 ng/ml TGF-β, both pseudopotentials at 1 ng/ml TGF-β have the new attractor closer to the epithelial state. Examination of individual trajectories also reveals that cells with 1 ng/ml TGF-β treatment leave the *E* region and mostly reside around a new attractor in the *I* region (Fig. S7 c&d). Some trajectories fluctuate with the attractor and only transiently reach the *M* region. That is, cells reach a new stable phenotype whose degree of mesenchymal features depends on the TGF-β concentration, which is consistent with previous studies on MCF10A cells treated with different concentrations of TGF-β ^13^. Fluctuations of single cell trajectories limit the state resolution in the cell feature space and prevent us from telling whether the two paths reach one common intermediate state or two distinct ones.

## DISCUSSION

Cell type regulation is an important topic in mathematical and systems biology. Recent advances of single cell techniques catalyzed quantitative studies on the dynamics of cell phenotypic transitions (CPT) emerging as a new field. How to study in the framework of dynamical systems theory and use experimental single cell techniques to compare with theoretical results. Here we present a quantitative framework for studying CPT dynamics using rate theories with time lapse imaging. In rate theories, identifying an appropriate RC provides mechanistic insights of a transition process. For example, recognizing collective solvent reorganization as part of the reaction coordinate is a key part of Marcus’s electron transfer theory ^29^. Adopting the fraction of native contacts as a RC also historically plays a key role in the development of protein folding theories. Therefore, continuous efforts have been made on defining an optimal RC for a complex dynamical system ^30, 31, 32^, and it is even questionable to assume that the system dynamics can be well presented by a 1-D RC, as suggested by theoretical analysis on protein folding ^33^. Indeed the analyses presented in this study show such breakdown of the basic assumption.

In introduction we discussed two mechanisms of critical state transitions. For TGF-β induced EMT in A549 cells, analyzing of the trajectories reveals that mathematically multiple paths may originate from destabilization of a multi-dimensional epithelial attractor through colliding with multiple saddle points sequentially. We illustrate such mechanism in Fig. 5f and in Movie S2 schematically with a metaphorical potential system. With no or low TGF-β, the system resides in the epithelial attractor (Fig. 5f, first). Adding TGF-β leads to appearance of a new attractor, and elevation of the epithelial attractor (Fig. 5f, second). The latter approaches and collides first with a saddle point to form a barrier-less (concerted-variation) path to the new attractor (Fig. 5f, third). At this TGF-β concentration (e. g., 1 ng/ml), some barrier still exists along an alternative (vimentin-first) path, which then disappears with further increase of TGF-β concentration again (as revealed in Fig. 5d) through saddle node collision (Fig. 5f, fourth). Indeed we observe the reactive trajectories predominantly assume the concerted-variation path under 1 ng/ml TGF-β treatment (Fig. S6a), while the vimentin-first path becomes dominant under 4 ng/ml TGF-β treatment (Fig. S6b).

A long-hold concept in cell and developmental biology is that cells exist as discrete and distinct phenotypes. Single cell genomics measurements have challenged such view and reveal that cells move along continuum manifolds ^34^. The live cell imaging studies presented here support such picture by demonstrating EMT as relaxation along a continuum manifold, as what also revealed in previous single cell RNA-seq studies ^8^. Compared to the snapshot studies, live cell trajectories provide further true information on the transition dynamics, and reveal that the EMT trajectories can be clustered as fluctuating around two distinct paths that connect the epithelial and mesenchymal phenotypes. The existence of two types of transition paths suggest that the previously proposed conceptual 1D EMT axis ^9^ is insufficient for understanding how EMT proceeds, and puts dynamics inferred indirectly from snapshot data questionable ^8^.

The concept of Waddington’s landscape has inspired numerous theoretical efforts of constructing a scalar potential for non-conservative systems. Cautious should be taken on applying such concept to examine the dynamics of a system, as external variables such as the TGF-β concentration actively modulate the dynamical landscape, e.g., attractors, and the scalar potentials only contain part of the dynamical information. The pseudo-potential defined in this work is only valid in the special case of projecting to a 1-D subspace, and ambiguity arises if one tries to connect the pseudo-potentials of the two convergent paths. Instead with more single cell trajectories one can reconstruct the multi-dimensional vector field directly following the procedure described in in this work.

In summary, through an integrated framework of live cell imaging and single cell trajectories with dynamical systems theories we obtained quantitative mechanistic insights of TGF-β induced EMT as a prototype for CPTs in general. Since a cell is a complex dynamical system with many strongly coupled degrees of freedom and a broad range of relevant time scales, further development will benefit from a finer resolution of cell state through including additional measurable cell features.

## METHODS

### 1) Time lapse imaging

A549/VIM-RFP (ATCC® CCL-185EMT™) were in F-12K medium (Corning) with 10% fetal bovine serum (FBS) in MatTek glass bottom culture dishes (P35G-0-10-C) in a humidified atmosphere at 37 □ and 5% CO2, as detailed in ^10^. Time-lapse images were taken with a Nikon Ti2-E microscope with differential interference contrast (DIC) and TRITC channels (Excitation wavelength is 555 nm and Emission wavelength is 587) (20 × objective, N.A. = 0.75). The cell culture condition was maintained with Tokai Hit Microscope Stage Top Incubator. For the 4 ng/ml TGF-β experiment, cells were imaged every 5 min with the DIC channel and every 10 min with the TRITC channel for two days. The exposure time for DIC was 100 millisecond (ms) and the exposure time for the TRITC channel was 30 ms. For the 1 ng/ml TGF-β experiment, cells were imaged every 5 min with the DIC channel and every 15 min with the TRITC channel for three days. The exposure time for DIC was 100 ms and the exposure time for the TRITC channel was 30 ms. While taking the images, all the imaging fields were chosen randomly.

### 2) Self-organizing map and shortest transition paths in the directed network

The self-organizing map is an unsupervised machine learning method to represent the topology structure of date sets. We used a 12 × 12 grid (neurons) to perform space approximation of all reactive trajectories. The SOM was trained for 50 epochs on the data with Neupy (*http://neupy.com/pages/home.html*). We set the learning radius as 1 and standard deviation 1. These neurons divide the data into 144 micro-clusters ({*ψ*}). With the single cell trajectory data, we counted the transition probabilities from cluster *i* to cluster *j* (including self), with 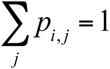. If the transition probability is smaller than 0.01, the value is then reset as 0. With the transition probability matrix, we built a directed network of these 144 neurons (Fig. S1). The distance (weight) of the edge between neuron *ψ*_*i*_ and neuron *ψ*_*j*_ is defined as the negative logarithm of transition probability (−log *p*_*i, j*_). We set the neurons that close to the center of epithelial and mesenchymal state (sphere with radius = 0.7) as epithelial community and mesenchymal community, respectively, and used Dijkstra algorithm to find the shortest path between each pair of epithelial and mesenchymal neurons ^20^ with NetworkX ^35^. We recorded the frequency of neurons and edge between these neurons that were past by these shortest paths.

### 3) Calculation of density of reactive trajectory

The density of reactive trajectory on the plane of morphology PC1 and vimentin Haralick PC1 was calculated with the following procedure:

a. Divide the whole plane into 200×200 grids.
b. In each grid, count the number of reactive trajectory (only the parts of each reactive trajectory that were in the intermediate region were taken into consideration) that enters and leaves it. If a reactive trajectory passes certain grid multiple times, only one was added in this grid’s density. Thus, the density matrix was obtained.
c. Use Gaussian filter to smooth the obtained density matrix. The standard deviation was set to be two and the truncation was two (i.e., truncate at twice of the standard deviation).

### 4) Procedure for determining a reaction coordinate

We followed a procedure adapted from what used in the finite temperature string method for numerical searching of reaction coordinate and non-equilibrium umbrella sampling ^23, 25^, with a major difference that we used experimentally measured single cell trajectories (Fig. S2).

a. Identify the starting and ending points of the reaction path as the means of data points in the epithelial and mesenchymal regions, respectively. The two points are fixed in the remaining iterations.
b. Construct an initial guess of the reaction path that connects the two ending points in the feature space through linear interpolation. Discretize the path with *N* (= 30) points (called images, and the *k*_*th*_ image denoted as *s*_*k*_ with corresponding coordinate **X**(*s*_*k*_)) uniformly spaced in arc length.
c. Collect all the reactive single cell trajectories that start from the epithelial region and end in the mesenchymal region.
d. For a given trial RC, divide the multi-dimensional state space by a set of Voronoi polyhedra containing individual images, and calculate the score function *F* given in the main text (with *w* = 10 in our calculations). We carried out the minimization procedure through an iterative process. For a given trial path defined by the set of image points, we calculated a set of average points using the following equations, 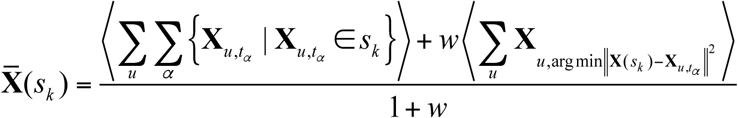. Next we updated the continuous reaction path through cubic spline interpolation of the average positions ^36^, and generated a new set of *N* images {*X* (*s*_*k*_)} that are uniformly distributed along the new reaction path. We set a smooth factor, *i.e*., the upper limit of 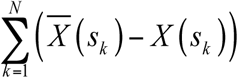, as 1 for calculating the RC in Fig. 4.
e. We iterated the whole process in step 3 until there was no further change of Voronoi polyhedron assignments of the data points.
f. For obtaining the pseudo-potential of a larger range of *s*, extrapolate the obtained reaction path forward and backward by adding additional image points (3 for the two parallel reaction paths) beyond the two ends of the path linearly, respectively. These new image points are also uniformly distributed along *s* as the old image sets do. Re-index the whole set of image points as {*s*_0_, *s*_1_,…, *s*_*i*_, …, *s*_*N*_, *s*_*N* +1_}.

### 5) Calculation of dynamics of morphology and Haralick features along reaction path

The reaction path is calculated in the principal component (PC) space of morphology PC1, vimentin Haralick PC1, PC3 and PC4. Distribution of cells show significant shift before and after TGF-β treatment in these dimensions ^10^. To reconstruct dynamics in the original features space from PCs, the reaction path’s coordinates on the other dimensions of PC are set as means of data in the corresponding Voronoi cell of each point on the reaction path. We obtain the reaction path in full dimension of PC space. The dynamics of morphology and Haralick features are calculated by inverse-transform of coordinates of PCs.

### 6) Markov property test

We examined the Markov property of the RC trajectories with Chapman-Kolmogorov Test (CK-test) (Fig. S3). The CK-test compares the left and right sides of Chapman-Kolmogorov equation (*P* (*kτ*) = *P*^*k*^(*τ*)). The *k*-step transition matrix should equal to the *k*_*th*_ power of 1-step transition matrix if the process is Markovian. We estimated the transition matrix with PyEMMA ^37^.

### 7) Reconstruction of pseudo-potential along the reaction coordinate

Based on the theoretical framework in Fig. 5c, we followed the procedure below:

a. The *N* + 2 image points of an identified RC divide the space in *N* + 2 Voronoi cells that data points can assign to. Ignore the first and last Voronoi cells, and use the remaining *N* cells for the remaining analyses.
b. Within the *i*_*th*_ Voronoi cell, calculate the mean drift speed (and thus *dϕ* / *ds*) at *s*_*i*_ approximately by 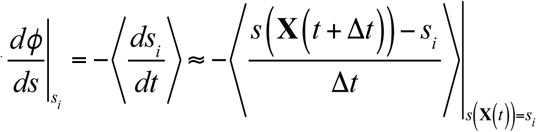 where *s*(**X**) is the assumed value along *s* for a cell state **X** in the morphology/texture feature space. The sum is over all time and all data points from all the recorded trajectories that lie within the *i*_*th*_ Voronoi cell (*s*(**X** (*t*)) = *s*_*i*_), and Δt =1 is one recording time interval. Using data points from all instead of just reactive trajectories is necessary for unbiased sampling within each Voronoi cell with 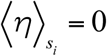.
c. Calculate the pseudo-potential through numerical integration, 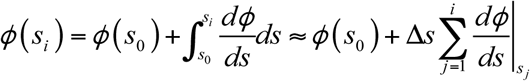. The exact value of ϕ(s_0_) does not affect the pseudo-potential shape.

### 8) Reconstruction of pseudo-potential along parallel paths

We followed the same mapping procedure for trajectories of cells treated with different TGF-β concentration. We used *tslearn* to calculate the DTW distance between two reactive trajectories, then performed K-Means clustering ^38^ on the DTW distance matrix ^39^ to cluster the reactive trajectories into two groups. We then followed the procedure in Methods (7) to reconstruct the RC for each group. We reconstructed the pseudo-potentials using all trajectories.

For a single cell trajectory, it is possible that the cell jumps out its original path due to fluctuation. For example, for a trajectory that mainly follows RC1, certain parts of it may transit into the range of RC2. So we used a part-aligning method to map all the trajectories to the RCs.

For a trajectory not belonging to the reactive trajectory ensemble, we assigned it to one of the two group associated to the two RCs, {*s*_1_} = {*s*_1,1_,…, *s*_1,*i*_, …, *s*_1,*N*_} and {*s*_2_} = {*s*_2,1_,…, *s*_2,*i*_, …, *s*_2,*N*_} by using sub-sequence DTW distance^38^. We first calculate the ub-sequence DTW distance of this trajectory to the Voronoi cells of {*s*_1_} and {*s*_2_}, respectively, and identified its matching coordinates {*s*_1,*a*_, …, *s*_1,*i*_, …, *s*_1,*b*_}, and {*s*_2,*c*_, …, *s*_2,*i*_, …, *s*_2,*d*_}on the RCs. Each point on the trajectory was assigned to the RCs based on minimum Euclidean distance. Then the consecutive parts of this trajectory along each RC were used to calculate the drift speed ⟨ *ds* / *dt* ⟩ and pseudo-potential *ϕ* (*s*) following the definition and procedure described in Methods (7).

### 9) Numerical solution of the Fokker-Planck equation

The FP equation can be transformed into Smoluchowski Diffusion equation 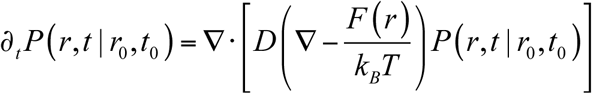. In the equation, *D*, which is spatially dependent, was estimate from the experiment data. 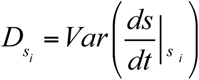 The variance of 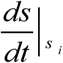 in corresponding Voronoi cells was calculated from experiment data. The term *F* (*r*) = −∇ *U*, where *U* is the pseudo-potential obtained from experiment data, the pseudo-temperature *T* was set as 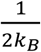, where *k*_*B*_ is the Boltzmann constant. With these inputs, we solved the equations numerically ^40^.

## Acknowledgments

This work was partially supported by National Cancer Institute (R37 CA232209), and National Institute of Diabetes and Digestive and Kidney Diseases (R01DK119232) to JX.

## Author Contributions

WW, JX conceived the project. WW, DP, YY and TH performed the experiment and data analysis. WW and JX wrote the manuscript.

**There is no competing interest**.

## Supplemental figures

**Figure S1.**
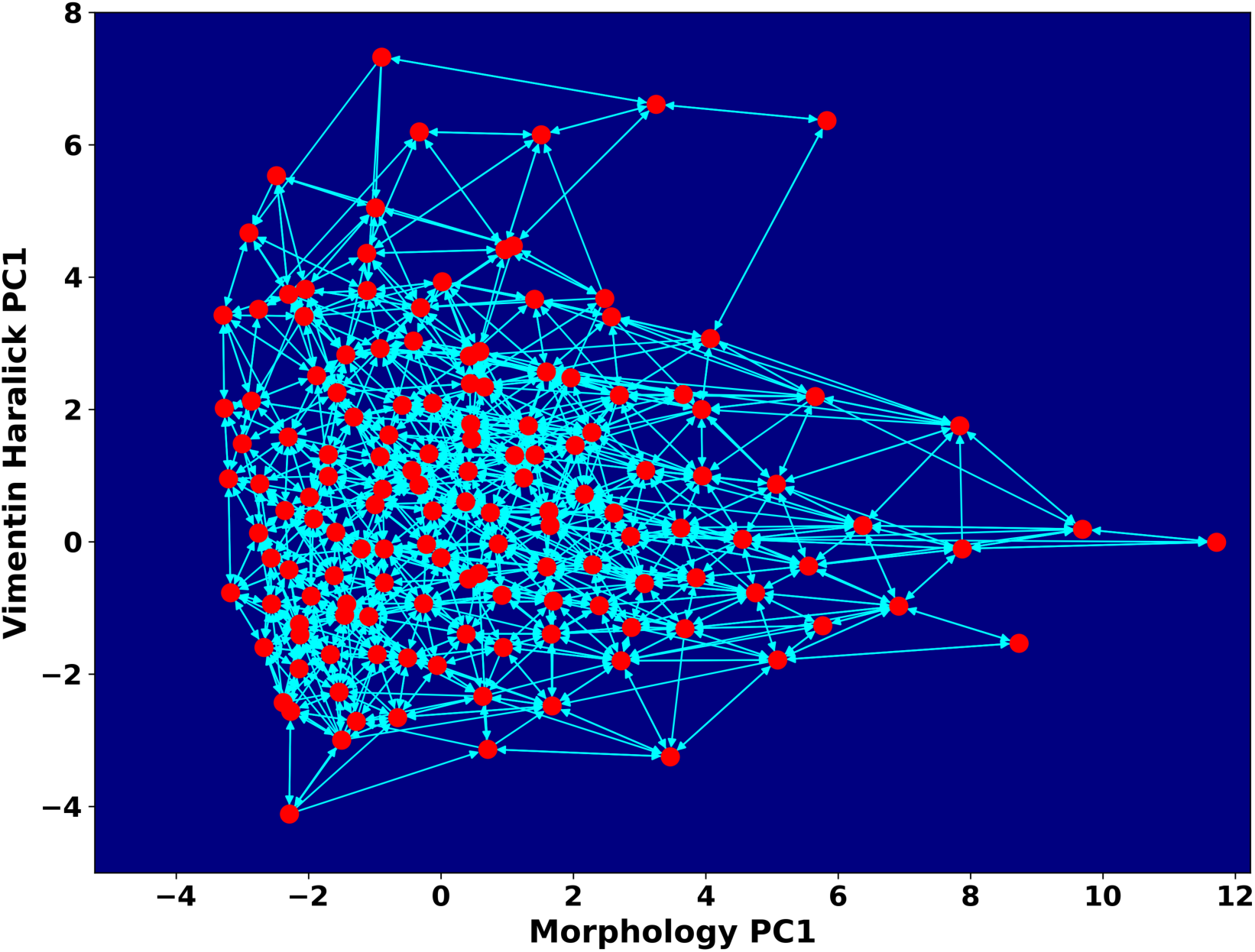
Space approximation of the whole single cell data set with self-organizing map into 12×12 discrete states (clusters). Directed network generated base on the self-organizing map and the transition between states. The distance between two states is defined as the negative logarithm of transition probability.

**Figure S2.**
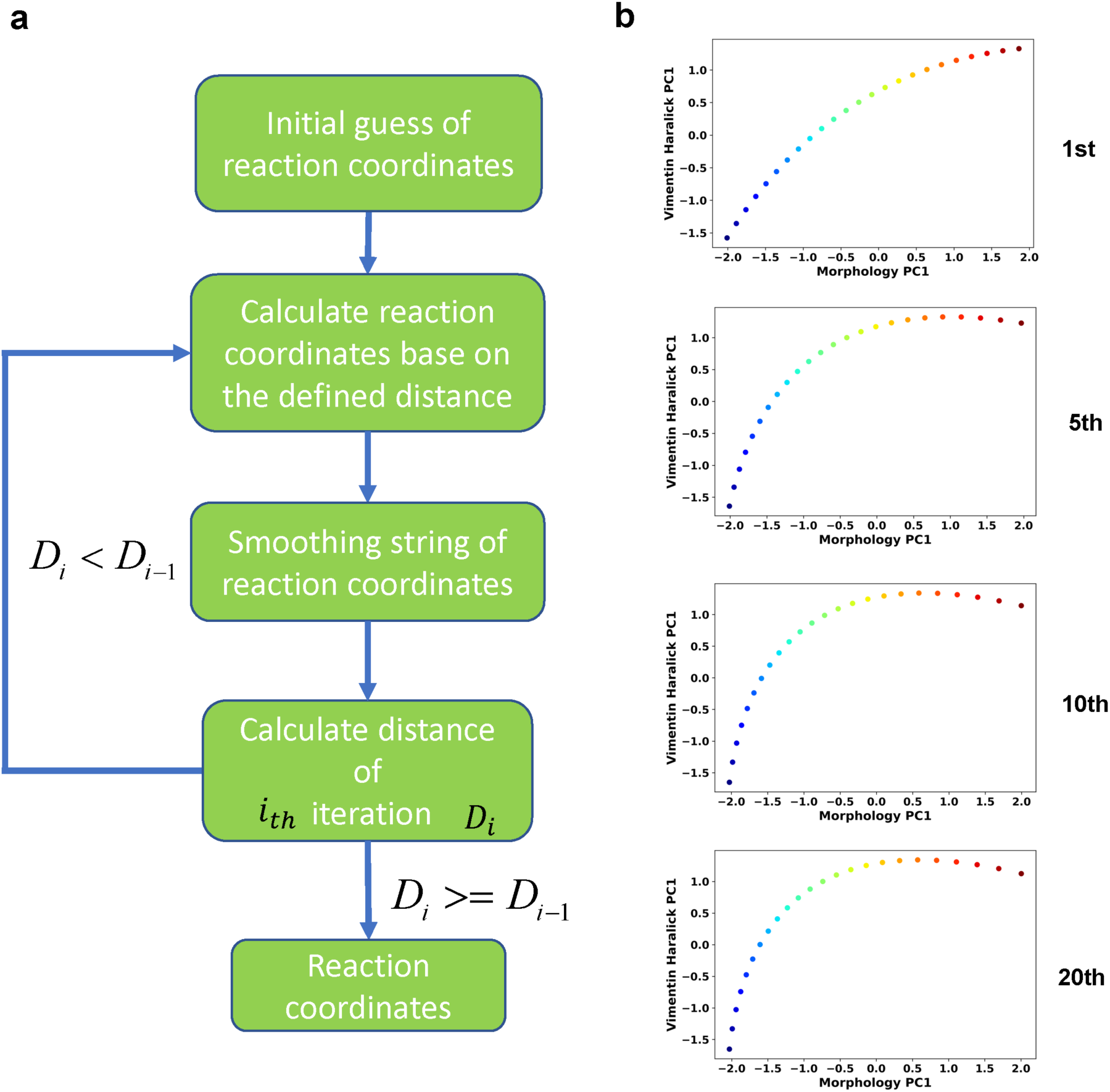
Iterative procedure of the finite temperature string method. **(a)** Flow chart of the procedure. (**b**) Example RC curves obtained at different iteration cycles

**Figure S3.**
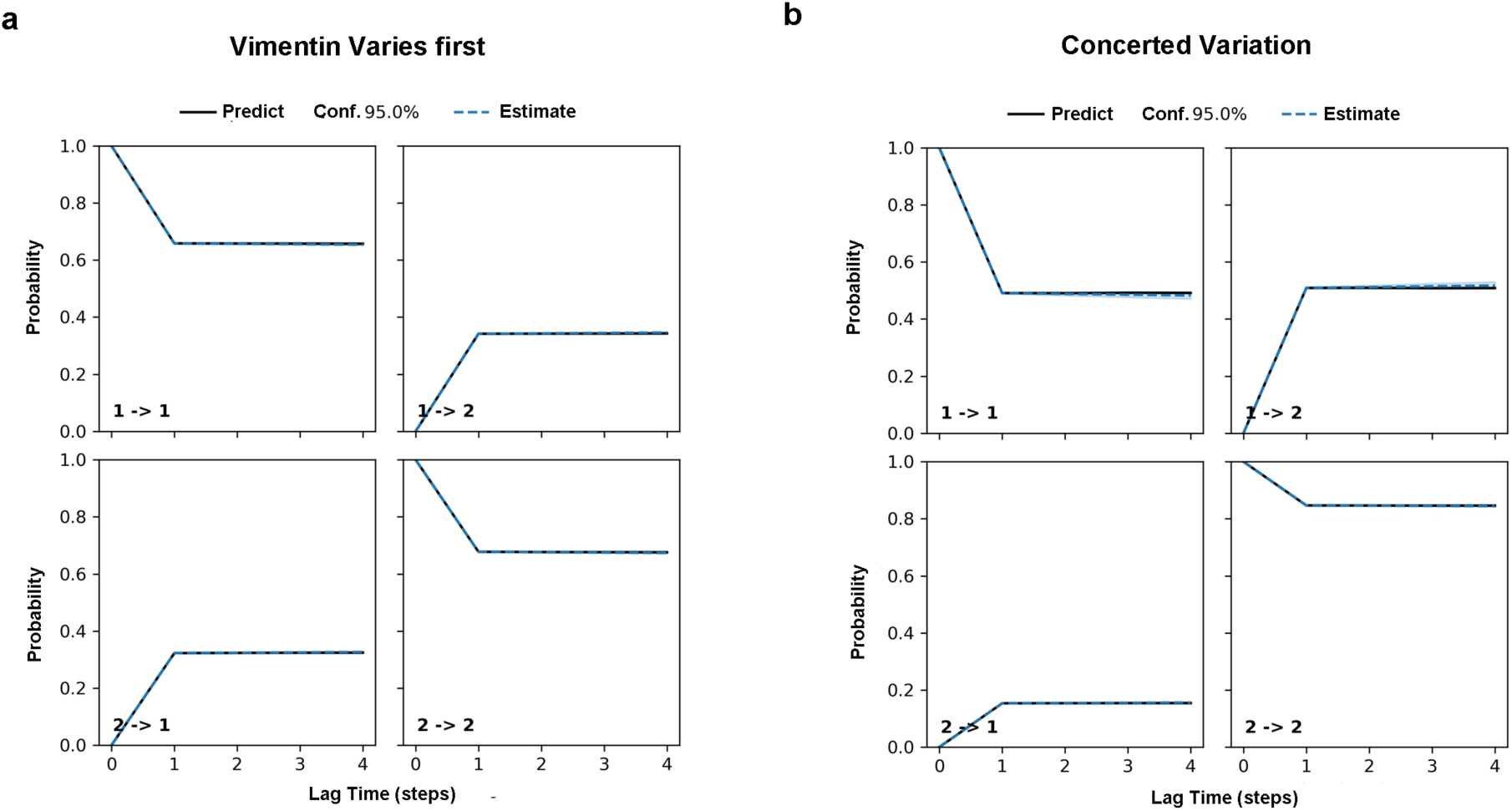
Chapman-Kolmogorow test on the RC s_1_ (a) and s_2_ (b). Black solid lines are model predictions estimated from one-step time lag. Blue dashed lines are the estimations from larger steps of time lag. The shading blue region is the region with 95% confidence.

**Figure S4.**
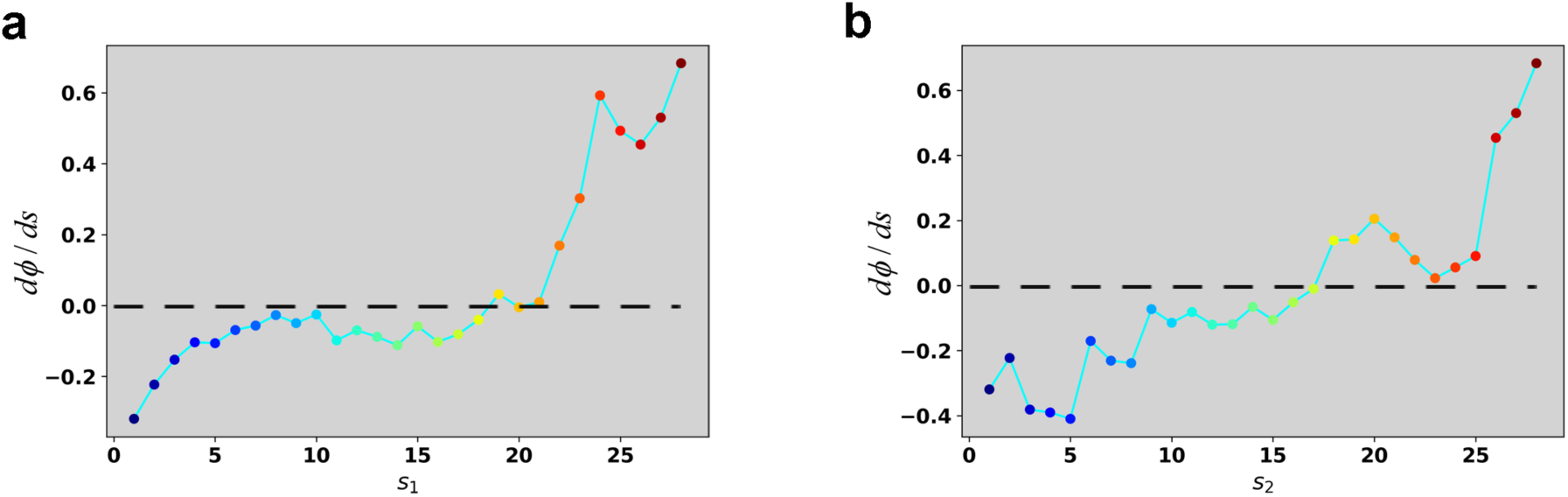
Reconstructed dϕ/ds along the RC in the case of 4 ng/ml TGF-β treatment. **(a)** Reconstructed dϕ/ds along the RC s_1._ **(b)** Reconstructed dϕ/ds along the RC s_2_.

**Figure S5.**
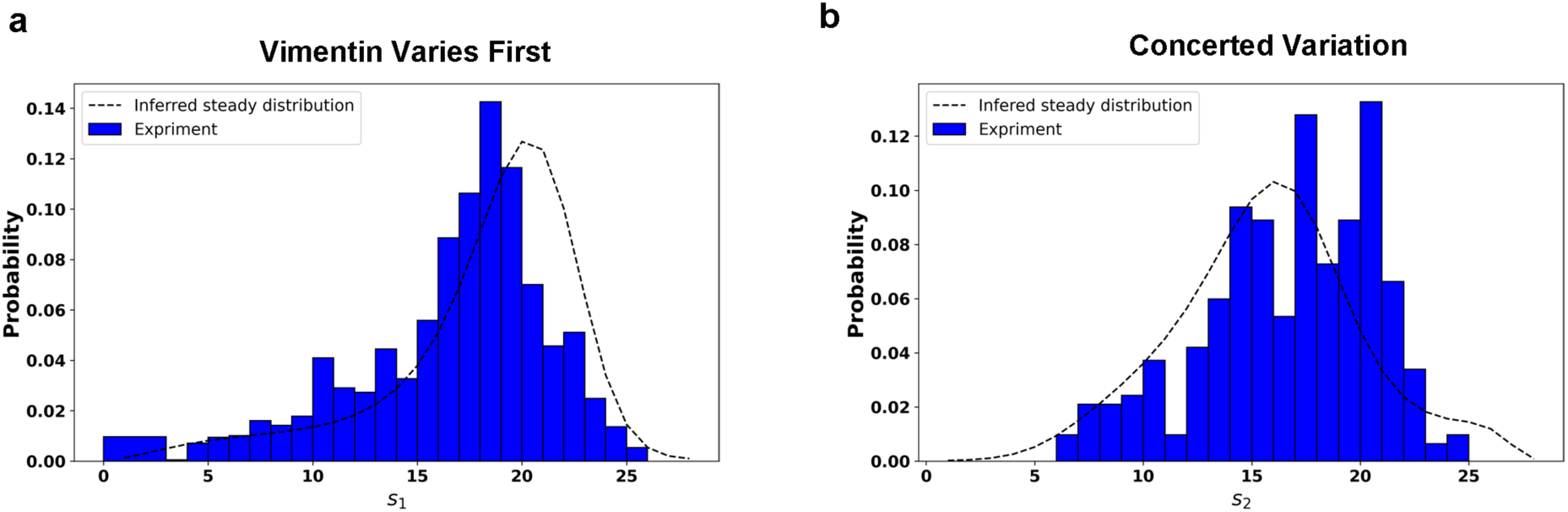
Comparison between the predicted steady distribution of RC values from reconstructed potential and experiment data along the RC s_1_ (a) and the RC s_2_ (b). The effective temperature value in calculation is set as 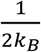.

**Figure S6.**
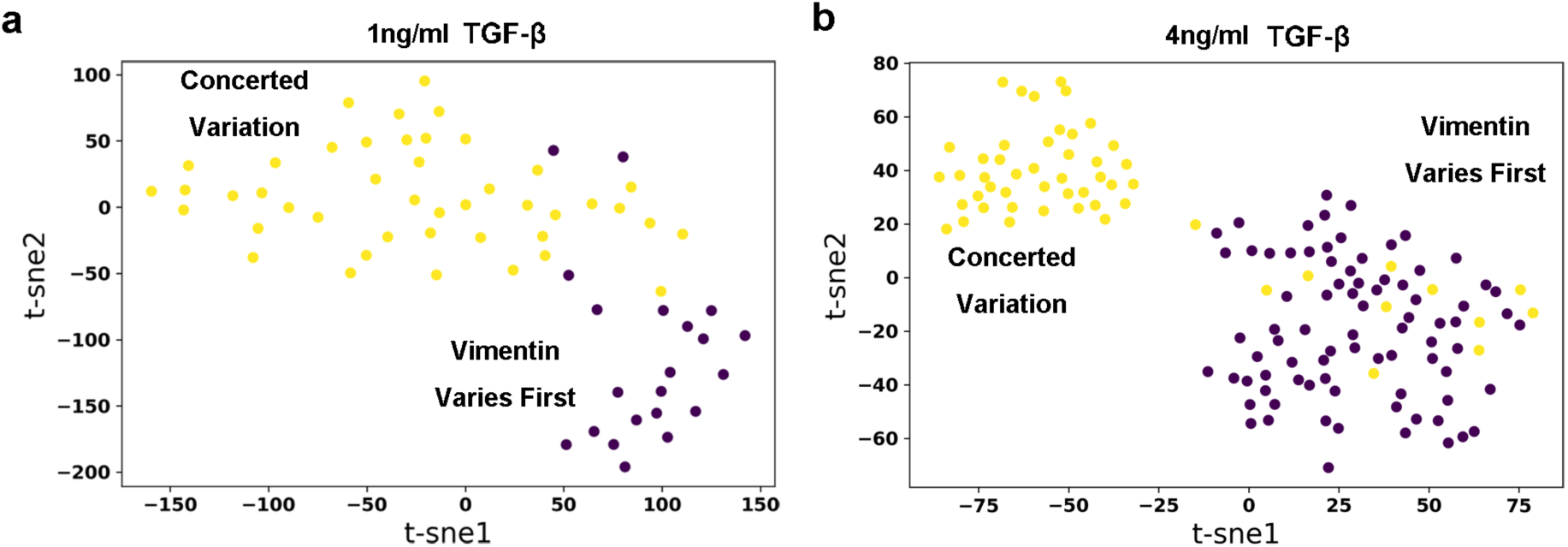
Embedding of dynamics time warping (DTW) distance matrix of reactive trajectories under 1 ng/ml (a) and 4 ng/ml (b) TGF-β treatment with t-SNE. Each dot represents a reactive trajectory. The colors of dots represent different clusters they belong to: Brown, vimentin varies first; Yellow, Concerted variation.

**Figure S7.**
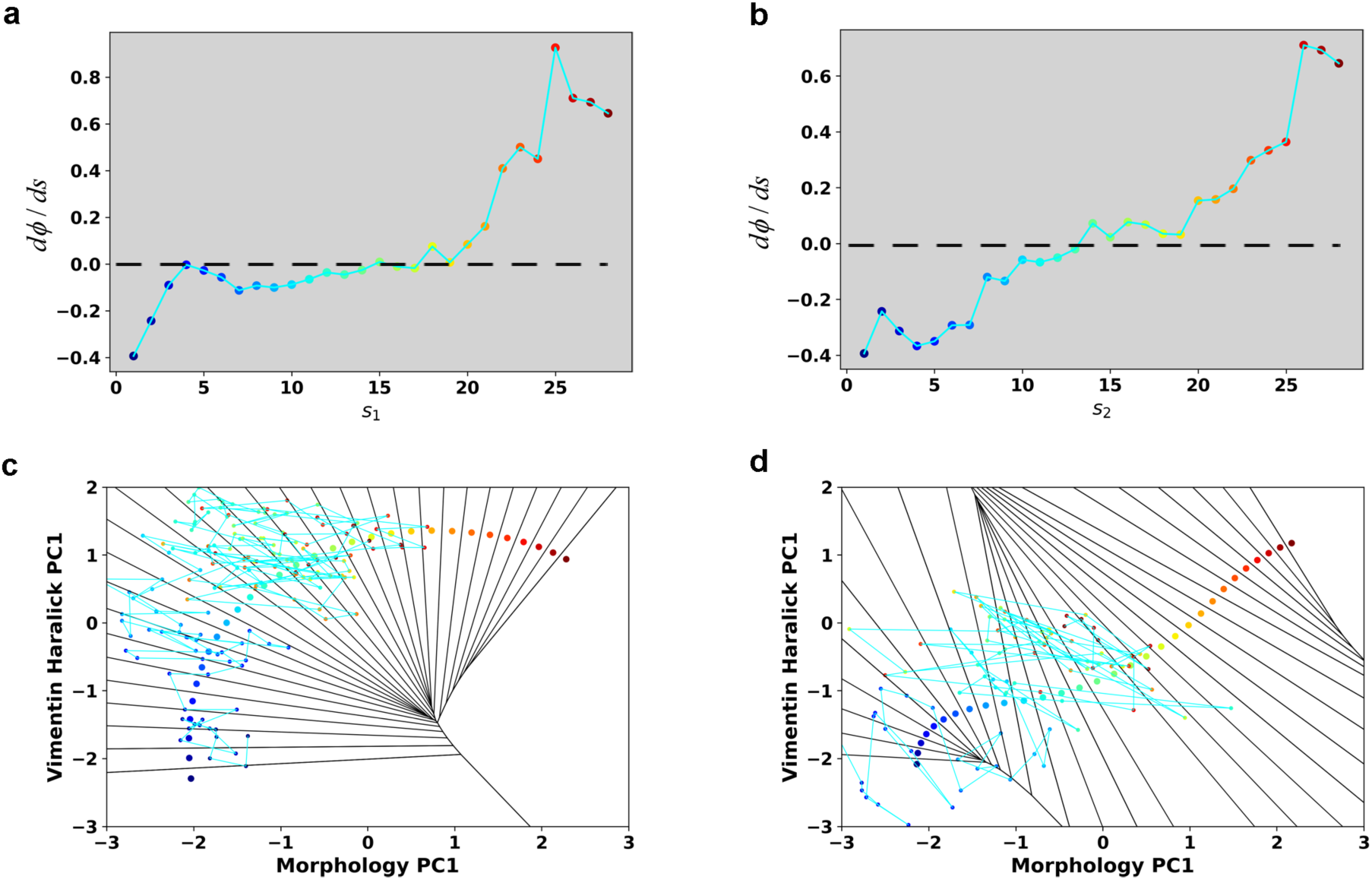
Additional analyses of trajectories with 1 ng/ml TGF-β treatment. **(a)** Reconstructed dϕ/ds along the RC s_1_ in the case of 1 ng/ml TGF-β treatment **(b)** Reconstructed dϕ/ds along the RC s_2_ in the case of 1 ng/ml TGF-β treatment. **(c)** Typical single cell trajectories treated with 1 ng/ml TGF-β along RC s_1_ **(d)** Typical single cell trajectories treated with 1 ng/ml TGF-β along RC s_2_

***Movie S1: Recorded live cell trajectory in Fig. 2e. Each frame is a segmented cell mask cropped from the original vimentin fluorescence image***.

***Movie S2: A metaphorical potential system illustrating how TGF-β treatment modifies the cell dynamics***.

## Notes

### Competing Interest Statement

The authors have declared no competing interest.

### Summary of Updates

The manuscript has been extensively revised with a shift of focus on identifying two parallel transition paths of EMT and a sequential saddle-node bifurcation mechanism.

## References

1. Wagner DE, Klein AM. Lineage tracing meets single-cell omics: opportunities and challenges. Nature Reviews Genetics, (2020).

2. Weinreb C, Rodriguez-Fraticelli A, Camargo FD, Klein AM. Lineage tracing on transcriptional landscapes links state to fate during differentiation. Science 367, eaaw3381 (2020).

3. Huang S, Eichler G, Bar-Yam Y, Ingber DE. Cell Fates as High-Dimensional Attractor States of a Complex Gene Regulatory Network. Phys Rev Lett 94, 128701 (2005).

4. Mojtahedi M, et al. Cell fate decision as high-dimensional critical state transition. PLoS biology 14, e2000640 (2016).

5. Chen L, Liu R, Liu Z-P, Li M, Aihara K. Detecting early-warning signals for sudden deterioration of complex diseases by dynamical network biomarkers. Sci Rep-Uk 2, (2012).

6. Tian XJ, Zhang H, Xing J. Coupled reversible and irreversible bistable switches underlying TGFβ-induced epithelial to mesenchymal transition. Biophys J 105, 1079–1089 (2013).

7. Yang J, et al. Guidelines and definitions for research on epithelial–mesenchymal transition. Nat Rev Mol Cell Biol, (2020).

8. McFaline-Figueroa JL, Hill AJ, Qiu X, Jackson D, Shendure J, Trapnell C. A pooled single-cell genetic screen identifies regulatory checkpoints in the continuum of the epithelial-to-mesenchymal transition. Nature Genetics 51, 1389–1398 (2019).

9. Nieto MA, Huang RY, Jackson RA, Thiery JP. Emt: 2016. Cell 166, 21–45 (2016).

10. Wang W, et al. Live-cell imaging and analysis reveal cell phenotypic transition dynamics inherently missing in snapshot data. Science Advances 6, eaba9319 (2020).

11. Ye Z, Sarkar CA. Towards a quantitative understanding of cell identity. Trends Cell Biol 28, 1030–1048 (2018).

12. Gordonov S, Hwang MK, Wells A, Gertler FB, Lauffenburger DA, Bathe M. Time series modeling of live-cell shape dynamics for image-based phenotypic profiling. Integr Biol (Camb) 8, 73–90 (2016).

13. Zhang J, et al. TGF-β-induced epithelial-to-mesenchymal transition proceeds through stepwise activation of multiple feedback loops. Science signaling 7, ra91 (2014).

14. Zhang J, et al. Spatial clustering and common regulatory elements correlate with coordinated gene expression. PLoS Comput Biol 15, e1006786 (2019).

15. Cootes TF, Taylor CJ, Cooper DH, Graham J. Active shape models-their training and application. Comput Vis Image Und 61, 38–59 (1995).

16. Cootes TF, Taylor CJ, Cooper DH, Graham J. Active shape models-their training and application. Computer vision and image understanding 61, 38–59 (1995).

17. Haralick RM. Statistical and structural approaches to texture. Proceedings of the IEEE 67, 786–804 (1979).

18. Hanggi P, Talkner P, Borkovec M. Reaction-rate theory: 50 years after Kramers. Rev Mod Phys 62, 254–341 (1990).

19. Kohonen T. Self-organized formation of topologically correct feature maps. Biological cybernetics 43, 59–69 (1982).

20. Dijkstra EW. A note on two problems in connexion with graphs. Numerische mathematik 1, 269–271 (1959).

21. Barrallo-Gimeno A, Nieto MA. The Snail genes as inducers of cell movement and survival: implications in development and cancer. Development 132, 3151–3161 (2005).

22. Rohrdanz MA, Zheng W, Clementi C. Discovering mountain passes via torchlight: methods for the definition of reaction coordinates and pathways in complex macromolecular reactions. Annu Rev Phys Chem 64, 295–316 (2013).

23. Vanden-Eijnden E, Venturoli M. Revisiting the finite temperature string method for the calculation of reaction tubes and free energies. The Journal of chemical physics 130, 05B605 (2009).

24. Allen RJ, Warren PB, ten Wolde PR. Sampling Rare Switching Events in Biochemical Networks. Phys Rev Lett 94, 018104 (2005).

25. Dickson A, Warmflash A, Dinner AR. Nonequilibrium umbrella sampling in spaces of many order parameters. The Journal of chemical physics 130, 02B605 (2009).

26. Mojtahedi M, et al. Cell Fate Decision as High-Dimensional Critical State Transition. PLOS Biology 14, e2000640 (2016).

27. Xing J. Mapping between dissipative and Hamiltonian systems. J Phys A: Math Theor 43, 375003 (2010).

28. Xing J, Kim KS. Application of the projection operator formalism to non-Hamiltonian dynamics. J Chem Phys 134, (2011).

29. Marcus RA. Electron transfer reactions in chemistry. Theory and experiment. Reviews of Modern Physics 65, 599–610 (1993).

30. Li W, Ma A. Recent developments in methods for identifying reaction coordinates. Molecular simulation 40, 784–793 (2014).

31. Best RB, Hummer G. Reaction coordinates and rates from transition paths. Proc Natl Acad Sci USA 102, 6732 (2005).

32. Fukui K. The Path of Chemical-Reactions - the Irc Approach. Acc Che Res 14, 363–368 (1981).

33. Udgaonkar JB. Multiple Routes and Structural Heterogeneity in Protein Folding. Annual Review of Biophysics 37, 489–510 (2008).

34. Wagner DE, Klein AM. Lineage tracing meets single-cell omics: opportunities and challenges. Nature Reviews Genetics 21, 410–427 (2020).

35. Hagberg A, Swart P, S Chult D. Exploring network structure, dynamics, and function using NetworkX. (ed^(eds). Los Alamos National Lab.(LANL), Los Alamos, NM (United States) (2008).

36. Jones E, Oliphant T, Peterson P. SciPy: Open source scientific tools for Python. (2001).

37. Scherer MK, et al. PyEMMA 2: A Software Package for Estimation, Validation, and Analysis of Markov Models. Journal of Chemical Theory and Computation 11, 5525–5542 (2015).

38. Tavenard R. tslearn: A machine learning toolkit dedicated to time-series data. (ed^(eds) (2017).

39. Sakoe H, Chiba S. Dynamic programming algorithm optimization for spoken word recognition. IEEE Transactions on Acoustics, Speech, and Signal Processing 26, 43–49 (1978).

40. Holubec V, Kroy K, Steffenoni S. Physically consistent numerical solver for time-dependent Fokker-Planck equations. Physical Review E 99, 032117 (2019).

